# Monitoring intramolecular dynamics across two regions of the mouse prion protein during misfolding and oligomerization using fluorescence correlation spectroscopy

**DOI:** 10.64898/2026.09.25.753467

**Authors:** Preeti Kumari

**Affiliations:** Department of Chemistry, Indian Institute of Science Education and Research Pune (IISER-Pune), Pune 411008, India

**Keywords:** Mouse prion protein (moPrP), Oligomerization, Protein aggregation, Prion misfolding, MEM, PET-FCS, single-molecule spectroscopy, Oligomer dynamics, Microsecond fluctuations

## Abstract

It is important to determine whether native state dynamics drive the misfolding and oligomerization of the prion protein, which are important events in prion disease, and how they are modulated by conformational conversion. Native (N) mouse prion protein (moPrP) is known to form small (O_S_) and large (O_L_) oligomers rich in β-sheet, and in this study, photoinduced electron transfer-fluorescence correlation spectroscopy (PET-FCS) has been used to characterize intramolecular dynamics within individual monomeric units in both isolated O_S_ and O_L_, as well as the diffusion properties of the oligomers. It is estimated that O_S_ and O_L_ comprise of about 15 and 55 monomeric units, respectively. Microsecond dynamics at each of the two regions that are the α1-α3 and α2-α3 interfaces of native protein are distinct in N, O_S_ and O_L,_ although they occur on very similar timescales. Analysis of the evolution of the distribution of diffusion times, determined using the maximum entropy method, indicates heterogeneity in the oligomerization reaction. Analysis of the change in the fluctuations that occur in two different timescales in the native state ensemble shows that they are damped more at the erstwhile α1-α3 interface than the erstwhile α2-α3 interface. The difference in the extent of damping at the erstwhile α1-α3 and α2-α3 interfaces can be explained on the basis of the structural changes known to occur across each region. The changes in dynamics occur concurrently in both regions, indicating that the structural changes accompanying conformational conversion also occur simultaneously during the oligomerization of moPrP.

## Introduction

Prion diseases, also known as transmissible spongiform encephalopathies, are fatal neurodegenerative disorders associated with the misfolding and aggregation of the mammalian prion protein. They include Creutzfeldt–Jakob disease, Gerstmann–Sträussler–Scheinker syndrome, fatal familial insomnia and kuru in humans, scrapie in sheep and goats, bovine spongiform encephalopathy in cattle, and chronic wasting disease in cervids (Prusiner, 1982, 1998; Hill & Collinge, 2000; Aguzzi & Polymenidou, 2004; Weissmann, 2004; Caughey & Baron, 2006). Prion diseases are associated with the conformational conversion of the cellular prion protein, PrP^C^, from a soluble, predominantly α-helical and protease-sensitive form into misfolded PrP^Sc^, which is multimeric, β-rich, aggregation-prone and relatively protease-resistant (Prusiner, 1982, 1998; Pan et al., 1993; Aguzzi & Polymenidou, 2004). The misfolding of PrP can give rise to different β-rich aggregated forms, including amyloid fibrils and soluble oligomers (Baskakov et al., 2002; Jain & Udgaonkar, 2011; Singh & Udgaonkar, 2015a).

Importantly, small non-fibrillar β-rich oligomers appear to be highly infectious and more neurotoxic than fibrils, making oligomer formation a particularly important process to understand (Silveira et al., 2005; Simoneau et al., 2007). PrP likely misfolds when it encounters acidic conditions in the endocytic/lysosomal pathway (Borchelt et al., 1992). *In vitro*, moPrP undergoes misfolding and oligomerization at acidic pH, with the protonation of His186 and/or Asp201 playing an important role in initiating the process (Singh & Udgaonkar, 2016). At pH 4, β-rich oligomers form readily, whereas at physiological pH, oligomerization is not experimentally accessible because only a very small fraction of PrP molecules would have these residues protonated (Singh & Udgaonkar, 2016). Importantly, the intrinsic propensity of PrP from different mammalian species to form β-structured oligomeric forms at pH 4 correlates with the susceptibility to prion disease (Khan et al., 2010). Hence, β-rich oligomers formed by moPrP at pH 4 serve as a useful model system for studying conformational events during prion misfolding.

The native prion protein consists of an intrinsically disordered N-terminal region (NTR) and a structured C-terminal domain (CTD) composed of three α-helices, α1, α2 and α3, and two short antiparallel β-strands, β1 and β2. The structured domain is stabilized by a disulfide bond linking α2 and α3, but despite its apparently well-defined fold, the native (N) state of PrP is unusually dynamic (Moulick & Udgaonkar, 2014). Native-state HDX studies have shown that the N state of moPrP exists in equilibrium with partially unfolded forms (PUFs) (Moulick et al., 2015). NMR studies have also shown that oligomerizing conditions perturb local interactions in the structured CTD, particularly in α2, α3 and the α2–α3 interface (Sengupta et al., 2017; Bhate et al., 2021). Structural studies of PrP oligomers and fibrils have shown that major conformational conversion occurs mainly in the CTD (Singh et al., 2012; Singh & Udgaonkar, 2013).

Native-state HDX-MS studies have shown that local structural fluctuations in moPrP play an important role in initiating misfolding. Pathogenic mutations and protonation of key residues increase local structural dynamics, especially in α1 and in the loop between α1 and β2, and these changes have been linked to faster oligomerization (Singh & Udgaonkar, 2015b, 2016). On the other hand, stabilization of α2 prevents oligomer formation, indicating that local stability within the structured CTD can control the misfolding reaction (Singh et al., 2014). Building on these observations, later HDX-MS studies showed that specific PUFs are linked directly to the initiation of misfolding. In particular, mutation of conserved Pro residues to Ala makes PUF2* more accessible from the N state. The apparent rate constant of misfolding correlates with the equilibrium population of PUF2*, suggesting that misfolding commences from it (Pal & Udgaonkar, 2022). Mutation of conserved aromatic residues further showed that misfolding can also commence from other PUFs, including PUF1 and PUF2**. Thus, PrP misfolding can initiate through multiple structurally distinct precursor conformations (Pal & Udgaonkar, 2023). Rigidifying the β2–α2 loop increases the energy gap between the N state and misfolding-prone PUFs, thereby reducing PUF accessibility and slowing down oligomer formation (Pal & Udgaonkar, 2024). These studies established that the PUFs are important precursors for misfolding and oligomerization. However, HDX-MS studies mainly identify the structural regions involved in the N ↔ PUF transitions and the equilibrium populations of the PUFs. It does not directly reveal the timescale of the relevant intramolecular dynamics, which fluctuations contribute to oligomerization, and how oligomer formation alters local dynamics within each monomeric unit. Hence, there is a need for a site-specific method that can monitor fast intramolecular dynamics during the conversion of monomeric PrP into β-rich oligomers.

Photoinduced electron transfer coupled to fluorescence correlation spectroscopy, PET-FCS, is a sensitive method for monitoring fast, site-specific conformational dynamics. In PET, the fluorescence of an oxazine fluorophore is quenched when it comes into van der Waals contact with an electron donor such as tryptophan, making the method highly sensitive to short-range structural fluctuations (Doose et al., 2009; Sauer & Neuweiler, 2014). When combined with FCS, these appear in the autocorrelation function (ACF), allowing local dynamics to be measured at very low protein concentrations, and over the nanosecond to millisecond timescales (Neuweiler et al., 2009; Sauer & Neuweiler, 2014). Thus, PET-FCS is well-suited for probing rapid intramolecular motions that may underlie conformational conversion in proteins.

The utility of PET-FCS to study protein fluctuations has been amply demonstrated in several protein systems. In the case of miniprotein, Trp-cage, it revealed microsecond folding kinetics and denatured-state flexibility under equilibrium conditions (Neuweiler et al., 2005). In the case of the BBL domain, folding appeared as a microsecond relaxation, and faster submicrosecond dynamics reported intrachain contact formation (Neuweiler et al., 2009). PET-FCS further allowed direct measurement of chain motions within a folding intermediate of engrailed homeodomain (Neuweiler et al., 2010). Nanosecond to microsecond domain motions were detected in the spider silk N-terminal domain, as well as the ionotropic glutamate receptor N-terminal domain (Jensen et al., 2011; Ries et al., 2014). More recently, PET-FCS was used to monitor microsecond dynamics during the binding-induced folding of tau-K18 (Sen et al., 2021). Thus, PET-FCS is well established as a sensitive method for detecting fast, site-specific dynamics in folded proteins, folding intermediates and intrinsically disordered proteins.

A previous PET-FCS study on moPrP used two site-specific probes to monitor microsecond conformational dynamics in the monomeric protein under native-like conditions at pH 4. The W144/C199-Atto construct was designed to monitor dynamics between α1 and α3, whereas the W171/C225-Atto construct was designed to monitor dynamics between α2 and α3 (Goluguri et al., 2019). These sites were chosen because it had been proposed that PrP misfolding involves separation of the β1–α1–β2 subdomain from the α2–α3 core (Eghiaian et al., 2007; Singh et al., 2014; Singh & Udgaonkar, 2015a, 2015b). Hence, fluctuations across the α1–α3 and α2–α3 regions could be relevant to misfolding. PET-FCS measurements showed fluctuations occurring over three well-separated sub-millisecond timescales. The two faster fluctuations occur within the N state ensemble. The fluorescent native-state, N, samples two PET-quenched conformations, N* and N**, through transient dye–Trp contact. The slowest fluctuation occurs between the unfolded state, U, and the folding intermediate, I. The addition of salt, which initiates the oligomerization of moPrP at pH 4, altered the timescale of fluctuations in the protein core, suggesting that N state dynamics are important for sampling aggregation-prone conformations (Goluguri et al., 2019).

In this PET-FCS study, the increase in hydrodynamic size during the oligomerization of moPrP has been characterized, and is shown to be accompanied by the damping of local microsecond dynamics. The fast PET-FCS amplitudes decreased at both the α1–α3 and α2–α3 interfaces, indicating reduced local conformational fluctuations during the formation of β-rich oligomers. The effect was slightly stronger at the α1–α3 interface, consistent with the proposed role of subdomain separation in PrP misfolding. Thus, the PET-FCS measurements link oligomer growth with site-specific dynamic changes during moPrP conformational conversion.

## Results

### Doping with labelled protein enabled PET-FCS measurements without altering the kinetics of misfolding

Our group previously developed a co-oligomerization strategy to study site-specific FRET during moPrP oligomerization, in which labelled dopant protein was mixed with excess Trp-less moPrP to suppress intermolecular FRET and isolate intramolecular conformational changes (Sengupta & Udgaonkar, 2019). A similar strategy is applied in the current work, where 10 nM labelled moPrP, either W144/C199-Atto moPrP or W171/C225-Atto moPrP, was used to dope 100 µM Trp-less moPrP (Figure 1). The W144/C199-Atto PET probe was used to monitor intramolecular dynamics at the α1–α3 interface, whereas the W171/C225-Atto PET probe was used to monitor dynamics at the α2–α3 interface within labelled moPrP molecules under oligomerizing conditions.

**Figure 1.**
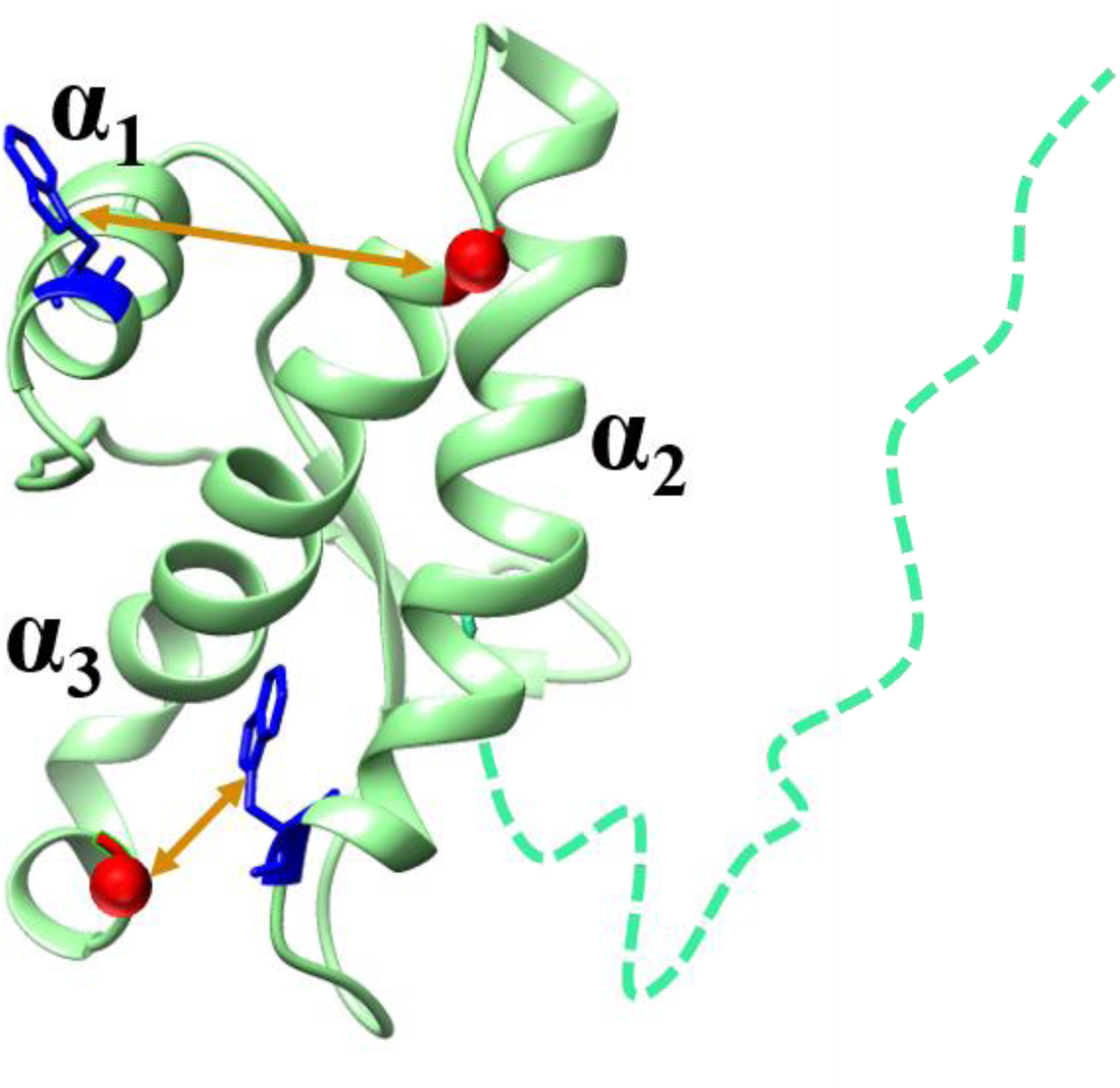
Structure of full-length moPrP highlighting the two PET pairs used to monitor dynamics across the two different subdomains and the core during misfolding and oligomerization. The W144/C199-Atto pair (top) was used to probe structural fluctuations between the α1 and α3 helices, representing dynamics across two different subdomains. The W171/C225-Atto pair (bottom) was used to monitor fluctuations within the α2-α3 core, reporting on dynamics within the same subdomain. Blue sticks represent the Trp residues which served as fluorescence quenchers, and red spheres indicate the Cys residues that were labelled with Atto 655. The brown arrows denote the distances across which dynamics were monitored. Protein Data Bank entry 1AG2 was used to make the figure using Chimera.

Before interpreting the PET-FCS measurements, it was necessary to establish that doping with labelled protein and the additives required for FCS did not perturb the oligomerization reaction. During oligomerization, wt moPrP is known to misfold and gain β-sheet structure at the expense of α-helical structure. Previous studies on wt moPrP had shown that the apparent rate constants of misfolding were the same whether monitored by far-UV CD at 222 nm, which is sensitive to α-helical structure, or at 216 nm, which is sensitive to β-sheet structure, and were similar to the rate constant of monomer loss during oligomerization (Singh et al., 2014; Sabareesan & Udgaonkar, 2016; Singh & Udgaonkar, 2016). In the present study, far-UV CD at 222 nm was used to monitor the loss of α-helical structure and the accompanying formation of β-rich oligomeric species. The far-UV CD-monitored kinetics of Trp-less moPrP doped with either W144/C199-Atto moPrP or W171/C225-Atto moPrP were similar to those reported earlier for Trp-less moPrP under the same oligomerization conditions (Figure 2) (Sengupta & Udgaonkar, 2019). Thus, doping with labelled protein did not alter the global misfolding reaction of Trp-less moPrP. For FCS measurements, 0.05% Tween 20 was added to minimize protein adsorption on the surface. Figure S1 shows that Tween 20 did not affect the kinetics of misfolding. Thus, neither doping with labelled moPrP nor the presence of 0.05% Tween 20 significantly altered the kinetics of moPrP misfolding under the conditions used here.

**Figure 2.**
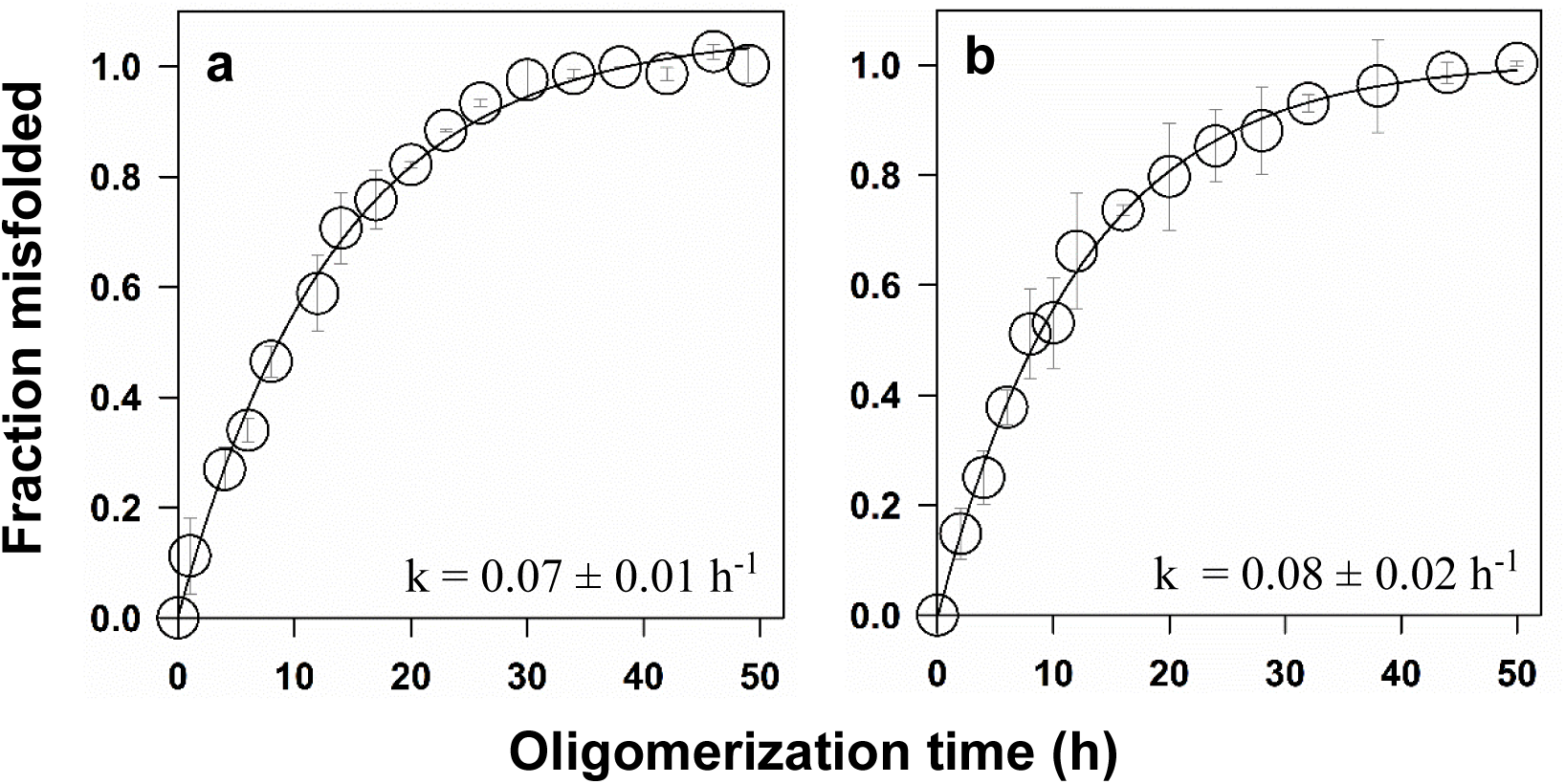
Kinetics of oligomerization of Trp-less moPrP doped with labelled moPrP. The conformational transition of 100 µM Trp-less mouse prion protein (moPrP) was monitored by far-UV CD at 222 nm. The reaction was carried out in the presence (a) 10 nM W144/C199-Atto moPrP and (b) 10 nM W171/C225-Atto moPrP. The solid lines represent fits of the data to a single exponential equation (Equation 1). The apparent rate constants obtained from the fits are shown in the panels. Oligomerization was carried in 150 mM NaCl, 10 mM NaOAc at pH 4.0 and 37 °C. The error bars represent standard deviations of the mean determined from two independent measurements.

### Oligomerization of Trp-less moPrP led to the formation of two oligomers, small and large

Figure S2a shows the size-exclusion chromatography (SEC) profile of the species present at the end of the oligomerization reaction. It was seen that at the end of the reaction, 8% of the protein was present as monomer (M), 43% as small oligomer (O_S_), and 49% as large oligomer (O_L_). Figure S2b shows the decrease in monomer concentration with time of oligomerization of 100 µM Trp-less protein. The rate constant for the disappearance of monomer was similar to the rate constant of the change in far-UV CD.

### FCS yielded the sizes of the small and large oligomers

To determine the diffusional properties of the species formed during oligomerization, the SEC-isolated monomeric, O_S_, and O_L_ fractions were and analyzed by PET-FCS. The ACFs, measured at 30 min and 60 min after isolation, were identical for both O_S_ and O_L_, indicating the interconversion between different oligomeric states did not occur over the timescale of the measurement. The ACFs provide information on the time taken (τ_D_) for the protein to diffuse through the confocal volume, and also on any faster dynamics that the protein may undergo as it traverses across the confocal volume. The ACFs of both monomers, W144/C199-Atto moPrP and W171/C225-Atto moPrP are similar in shape to those reported in an earlier study (Goluguri et al., 2019). When the ACFs of the monomers were analyzed using Equation 2, which portrays an ACF with one diffusion component (τ_D_) and three exponential processes describing faster fluctuations of the PET pairs, the values of the kinetic parameters that were obtained (Table 1) were similar to those reported earlier (Goluguri et al., 2019) (Figure S3; Table S1).

**Table 1.** Parameters obtained from analysis of ACFs of the monomer, small and large oligomers under oligomerization conditions.

| **W144/C199-Atto moPrP** |  |  |  |
| --- | --- | --- | --- |
| **Parameters** | **M** | **O<sub>s</sub>** | **O<sub>L</sub>** |
| **K<sub>1</sub>** | 0.9 ± 0.4 | 0.55 ± 0.02 | 0.30 ± 0.06 |
| **τ<sub>1</sub> (μs)** | 0.6 ± 0.4 | 0.31 ± 0.02 | 0.37 ± 0.03 |
| **K<sub>2</sub>** | 1.4 ± 0.3 | 0.13 ± 0.01 | 0.60 ± 0.03 |
| **τ<sub>2</sub> (μs)** | 2.8 ± 1.0 | 3.4 ± 0.15 | 2.5 ± 0.04 |
| **K<sub>3</sub>** | 0.14 ± 0.03 | 0.05 ± 0.004 | 0.10 ± 0.01 |
| **τ<sub>3</sub> (μs)** | 67 ± 4 | 34 ± 3 | 37 ± 2 |
| **τ<sub>D</sub> (μs)** | 283 ± 9 | 640 ± 28 | 948 ± 29 |
| **R<sub>h</sub> (nm)** | 2.75 ± 0.09 | 6.22 ± 0.27 | 9.21 ± 0.28 |
| **W171/C225-Atto moPrP** |  |  |  |
| **Parameters** | **M** | **O<sub>s</sub>** | **O<sub>L</sub>** |
| **K<sub>1</sub>** | 0.8 ± 0.05 | 0.53 ± 0.02 | 0.52 ± 0.004 |
| **τ<sub>1</sub> (μs)** | 0.4 ± 0.03 | 0.35 ± 0.04 | 0.6 ± 0.03 |
| **K<sub>2</sub>** | 0.5 ± 0.01 | 0.26 ± 0.01 | 0.30 ± 0.02 |
| **τ<sub>2</sub> (μs)** | 3 ± 0.3 | 3.0 ± 0.3 | 3.7 ± 0.5 |
| **K<sub>3</sub>** | 0.08 ± 0.01 | 0.11 ± 0.02 | 0.10 ± 0.01 |
| **τ<sub>3</sub> (μs)** | 34.5 ± 0.5 | 31 ± 6 | 39 ± 8 |
| **τ<sub>D</sub> (μs)** | 277 ± 8 | 752 ± 9 | 1187 ± 23 |
| **R<sub>h</sub> (nm)** | 2.69 ± 0.08 | 7.31 ± 0.09 | 11.54 ± 0.22 |

K<sub>1</sub>, K<sub>2</sub> and K<sub>3</sub> are the amplitudes and τ<sub>1</sub>, τ<sub>2</sub> and τ<sub>3</sub> are the time constants of the microsecond fluctuations. τ<sub>D</sub> is the translational diffusion time. Measurements were carried out in 150 mM NaCl, 10 mM NaOAc, pH 4.0 at 37 °C. The errors represent the standard deviations obtained from two independent experiments. R<sub>h</sub> is hydrodynamic radius obtained using the Stokes–Einstein equation.

The ACFs of the isolated oligomers showed the expected shift to longer diffusion times with increasing oligomer size (Figure 3). The diffusion times obtained by fitting the ACFs of the oligomers are listed in Table 1. For the monomer, as well as for the oligomers, the values of the diffusion times were used to determine the hydrodynamic radii, R_h_ (see Methods), which are also listed in Table 1. With the assumption that the monomer and both oligomers are spherical with similar partial molar volume, the number of monomeric units was determined to be 15 in O_S_ and 55 in O_L_ (see Methods).

**Figure 3.**
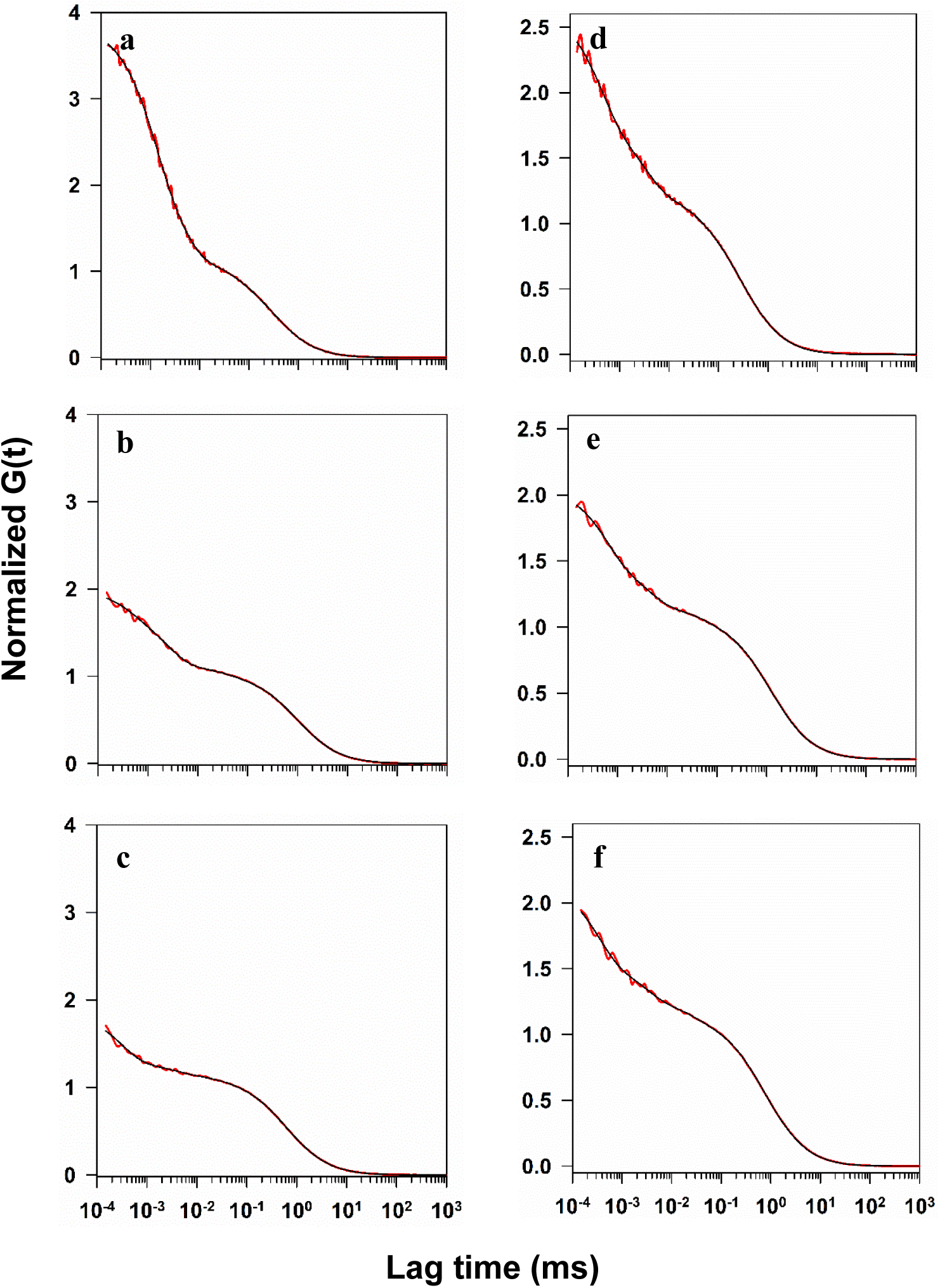
Conformational dynamics in the monomer, small and large oligomers in aggregation conditions: Panels a, b and c show the ACFs of the monomer, large oligomer and small oligomer, respectively of W144/C199-Atto moPrP doped Trp-less moPrP. Panels d, e and f show the ACFs of the monomer, large oligomer and small oligomer, respectively of W171/C225-Atto moPrP doped Trp-less moPrP. The ACFs were acquired on SEC purified samples in aggregation buffer (150 mM NaCl, 10 mM NaOAc, pH 4.0) at 37 °C. The red line in each panel is the data, and the black line is a fit to Equation 2. The value of the parameters obtained from each fit are listed in Table 1.

### Microsecond dynamics were modulated differentially in the large and small oligomers

The ACFs also provided kinetic data on the three microsecond conformational fluctuations occurring in monomeric units present in both the small and large oligomers (Figure 3). The time constants τ_1_, τ_2_ and τ_3_ of the three exponential processes and their respective amplitudes K_1_, K_2_ and K_3_ are listed in Table 1. When the dynamics were monitored using the W144/C199-Atto probe, the amplitudes of all three fast fluctuations were reduced for monomeric units present in both O_S_ and O_L_ with respect to the native monomer. However, the reduction was not identical in O_S_ and O_L_: the largest reduction in K_1_ was observed in O_L_, whereas K_2_ and K_3_ were reduced relatively more in O_S_. The corresponding time constants showed more limited changes; τ_1_ and τ_3_ were shorter in both O_S_ and O_L_, while τ_2_ was largely unaffected.

When the dynamics were monitored using the W171/C225-Atto probe, K_1_ and K_2_ were reduced in the monomeric units of both O_S_ and O_L_ to similar extents, whereas K_3_ was not detectably altered. τ_1_ was found to have increased only in O_L_ but not in O_S_, while τ_2_ and τ_3_ remained largely unchanged. Thus, oligomer formation did not abolish microsecond dynamics within a monomeric unit in either oligomer. But the amplitudes and, in a more limited manner, the time constants of the motions detected at the two sites were modified, in a different way in O_S_ and in O_L_.

### Evolution of PET-FCS autocorrelation functions during oligomerization

PET-FCS measurements were made at different times during the process of oligomerization to monitor the evolution of the size of the oligomer and local dynamics. Figure 4 shows the ACFs obtained at different times of oligomerization. The ACFs show that oligomerization had a bigger effect on the dynamics measured by the W144/C199-Atto PET pair than on those monitored using W171/C225-Atto PET pair. However, it was not possible to resolve the three time constants of diffusion in the diffusion component of the ACF, which may arise from M, O_S_ and O_L_ present at each time of oligomerization. Hence, at each time of oligomerization, the diffusion component of the ACF was fit to a single time constant, which was the weighted average of the time constants of diffusion of M, O_S_ and O_L_. Morever, the microsecond dynamics arising from each of M, O_S_, and O_L_ were not resolved from the above data, and hence conformational dynamics were fit to three exponentials, so that each time constant was a composite weighted average of the dynamics in each of M, O_S_ and O_L_. Table 2 gives the parameters obtained from the fits carried out in this manner according to Equation 2, which show a significant change during the process of oligomerization. In this context, it is important to note that only 1 in 10,000 of the oligomerizing protein molecules is a labelled dopant protein molecule. Hence, the likelihood of any oligomer that formed, whether O_S_ with 15 monomeric units or O_L_ with 55 monomeric units (see above), contained more than one labelled dopant protein molecule was negligible. Hence, the brightness of M, O_S_ and O_L_ would be similar.

**Figure 4.**
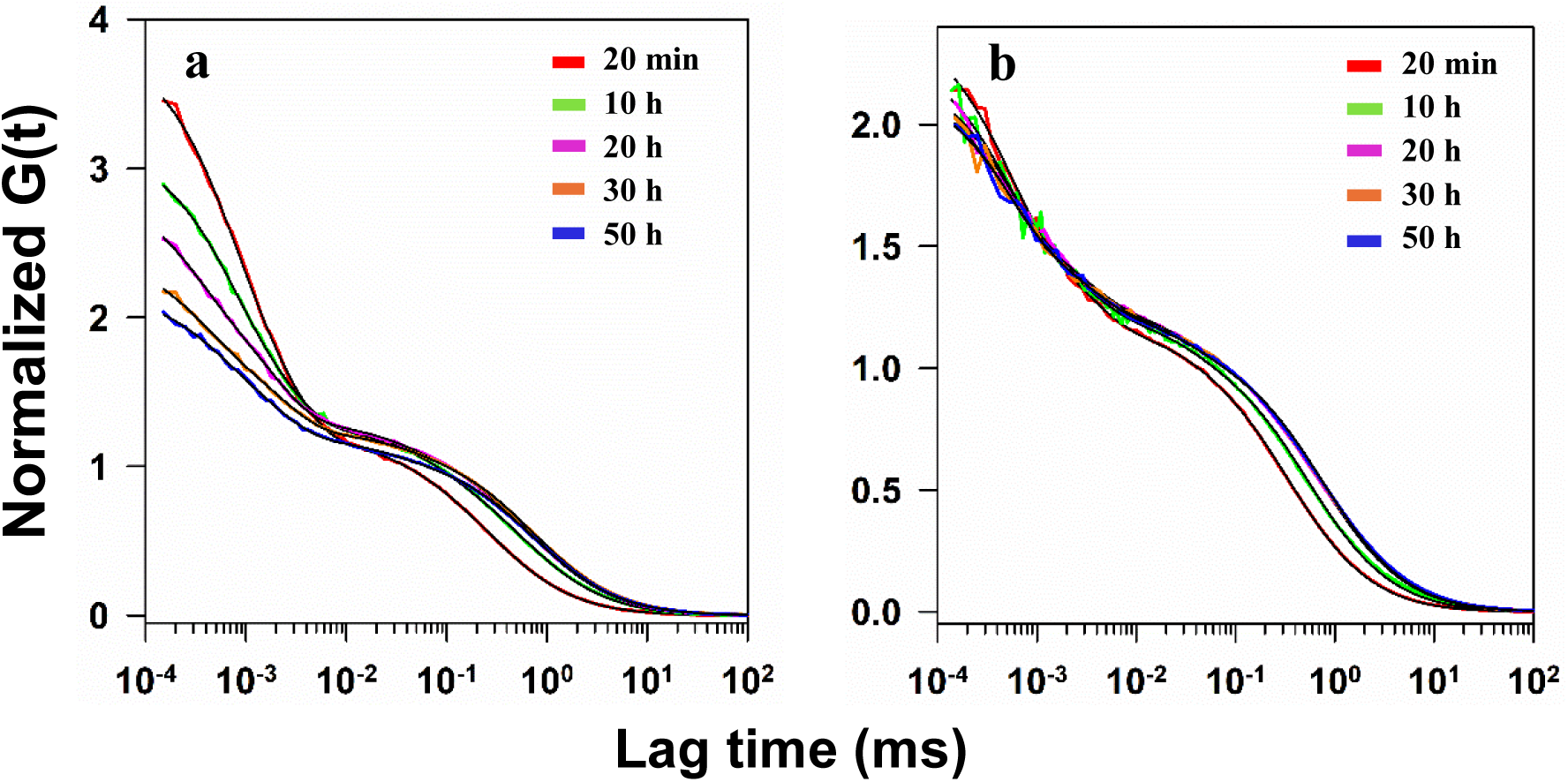
Evolution of microsecond conformational dynamics and diffusion during the oligomerization of moPrP in 150 mM NaCl, 10 mM NaOAc, pH 4.0 at 37 °C. Normalized ACFs are shown for 100 µM Trp-less moPrP doped with (a) 10 nM W144/C199-Atto moPrP and (b) 10 nM W171/C225-Atto moPrP at specific times of oligomerization. The black solid lines through the data represent fits to Equation 2.

**Table 2.** Parameters describing the ACFs obtained at different times of oligomerization.

| **W144/C199-Atto moPrP** |  |  |  |  |  |
| --- | --- | --- | --- | --- | --- |
| **Parameters** | **20 min** | **10 h** | **20 h** | **30 h** | **50 h** |
| **K<sub>1</sub>** | 1.1 ± 0.09 | 0.77 ± 0.11 | 0.59 ± 0.06 | 0.47 ± 0.009 | 0.45 ± 0.003 |
| **K<sub>2</sub>** | 1.2 ± 0.04 | 0.78 ± 0.06 | 0.58 ± 0.07 | 0.45 ± 0.003 | 0.44 ± 0.01 |
| **K<sub>3</sub>** | 0.07 ± 0.01 | 0.16 ± 0.01 | 0.12 ± 0.002 | 0.11 ± 0.001 | 0.11 ± 0.002 |
| **τ<sub>1</sub> (μs)** | 0.5 ± 0.08 | 0.52 ± 0.05 | 0.42 ± 0.01 | 0.5 ± 0.07 | 0.51 ± 0.05 |
| **τ<sub>2</sub> (μs)** | 2.2 ± 0.1 | 2.22 ± 0.08 | 2.28 ± 0.06 | 2.19 ± 0.01 | 2.265 ± 0.004 |
| **τ<sub>3</sub> (μs)** | 34 ± 2.9 | 58 ± 9.9 | 41 ± 5.9 | 38 ± 6.3 | 40 ± 5.4 |
| **τ<sub>D</sub> (μs)** | 307 ± 62 | 547 ± 12 | 639 ± 8 | 720 ± 1 | 734 ± 5 |
| **W171/C225-Atto moPrP** |  |  |  |  |  |
| **Parameters** | **20 min** | **10 h** | **20 h** | **30 h** | **50 h** |
| **K<sub>1</sub>** | 0.75 ± 0.003 | 0.62 ± 0.02 | 0.5 ± 0.05 | 0.52 ± 0.07 | 0.51 ± 0.05 |
| **K<sub>2</sub>** | 0.4 ± 0.02 | 0.3 ± 0.02 | 0.27 ± 0.05 | 0.3 ± 0.04 | 0.27 ± 0.1 |
| **K<sub>3</sub>** | 0.07 ± 0.0004 | 0.14 ± 0.02 | 0.14 ± 0.03 | 0.14 ± 0.02 | 0.14 ± 0.01 |
| **τ<sub>1</sub> (μs)** | 0.46 ± 0.02 | 0.46 ± 0.03 | 0.46 ± 0.01 | 0.46 ± 0.01 | 0.48 ± 0.02 |
| **τ<sub>2</sub> (μs)** | 2.8 ± 0.15 | 2.7 ± 0.1 | 2.8 ± 0.05 | 2.75 ± 0.06 | 2.7 ± 0.2 |
| **τ<sub>3</sub> (μs)** | 29 ± 1 | 30 ± 0.6 | 33 ± 0.01 | 30 ± 2 | 32 ± 1.7 |
| **τ<sub>D</sub> (μs)** | 327 ± 10 | 574 ± 63 | 683 ± 42 | 752 ± 28 | 790 ± 58 |

### Evolution of size during oligomerization

Figure 5 shows how the composite diffusional dynamics of the three species, M, O_S_ and O_L_, changed during oligomerization. Figures 5a and d show that the rate constant of the change in the composite diffusion time, τ_D_, was the same as that for the change in far-UV CD at 222 nm. The increase in the apparent diffusion time during the course of oligomerization for proteins doped with either labelled protein, show the formation of slow diffusing larger species with time. Since the time constants of diffusion of M, O_S_ and O_L_ could not be resolved using discrete analysis of the ACFs, the ACF at each time of oligomerization was analyzed using the model-free maximum entropy method (MEM) (see Methods). Figures 5b and e show the distribution of diffusion times, corresponding to τ_D_, at each time of oligomerization, which were obtained from MEM analysis. The distributions at different times of oligomerization were broader than that seen for the monomer, but the distribution of diffusion times at each time of oligomerization could not be resolved into distributions arising from M, O_S_ and O_L_. Figures 5c and f show that the peaks of the MEM distributions shifted with time of oligomerization with apparent rate constants similar to the rate constants of the change in far-UV CD.

**Figure 5.**
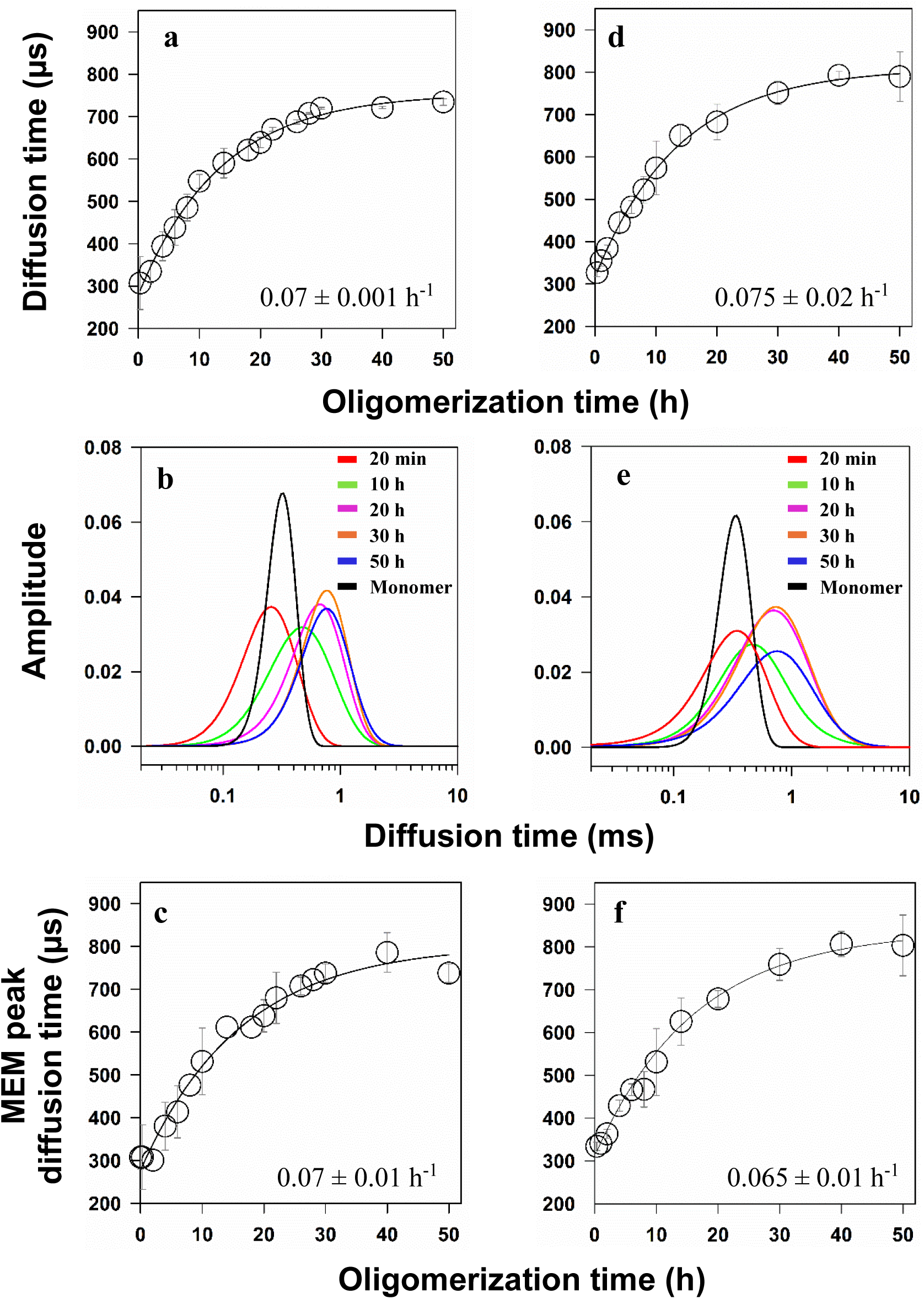
Evolution of diffusional dynamics during the oligomerization of moPrP. Oligomerization of 100 µM Trp-less moPrP doped with (a-c): W144/C199-Atto moPrP and (d-f): W171-C225Atto moPrP was carried out in aggregation buffer. (a,d) Evolution of the diffusion time (τ_D_): The increase in the average diffusion time determined from single diffusion component fits to Equation 2 as a function of oligomerization time. The apparent rate constants obtained from single exponential fits (solid lines through the data) are shown in the panels. (b,e) MEM distributions of diffusion times at different time of oligomerization, as well as of monomer in 10 mM NaOAc at pH 4. (c,f) MEM peak diffusion time evolution, time-dependent shift of the peak position of the MEM distributions. The apparent rate constants obtained from single exponential fits (solid lines through the data) are shown in the panels.

### Evolution of microsecond dynamics during oligomerization followed similar kinetics at the two sites

Figure 6 shows how the amplitudes of the fluctuations over three different timescales changed with oligomerization time at the two monitored sites. Figure S4 shows the dependence of the time constants of the three fluctuations on the time of oligomerization. At both the α1-α3 interface monitored by the W144/C199-Atto PET pair and the α2-α3 interface monitored by the W171/C225-Atto PET pair, the amplitudes of the fluctuations, K_1_ and K_2_, decreased with a rate constant similar to that of the change in far-UV CD. The time constants of the fluctuations remained invariant at both the interfaces. At both the interfaces, the amplitude K_3_ shows an increase in amplitude, which is much faster than the change in K_1_ and K_2_. The decreases in K_1_ and K_2_ indicated that, as the reaction progresses, the equilibrium shifts toward the bright state. As the oligomerization of the protein initiates with subdomain separation, it expected that the population of the dark state will be reduced.

**Figure 6.**
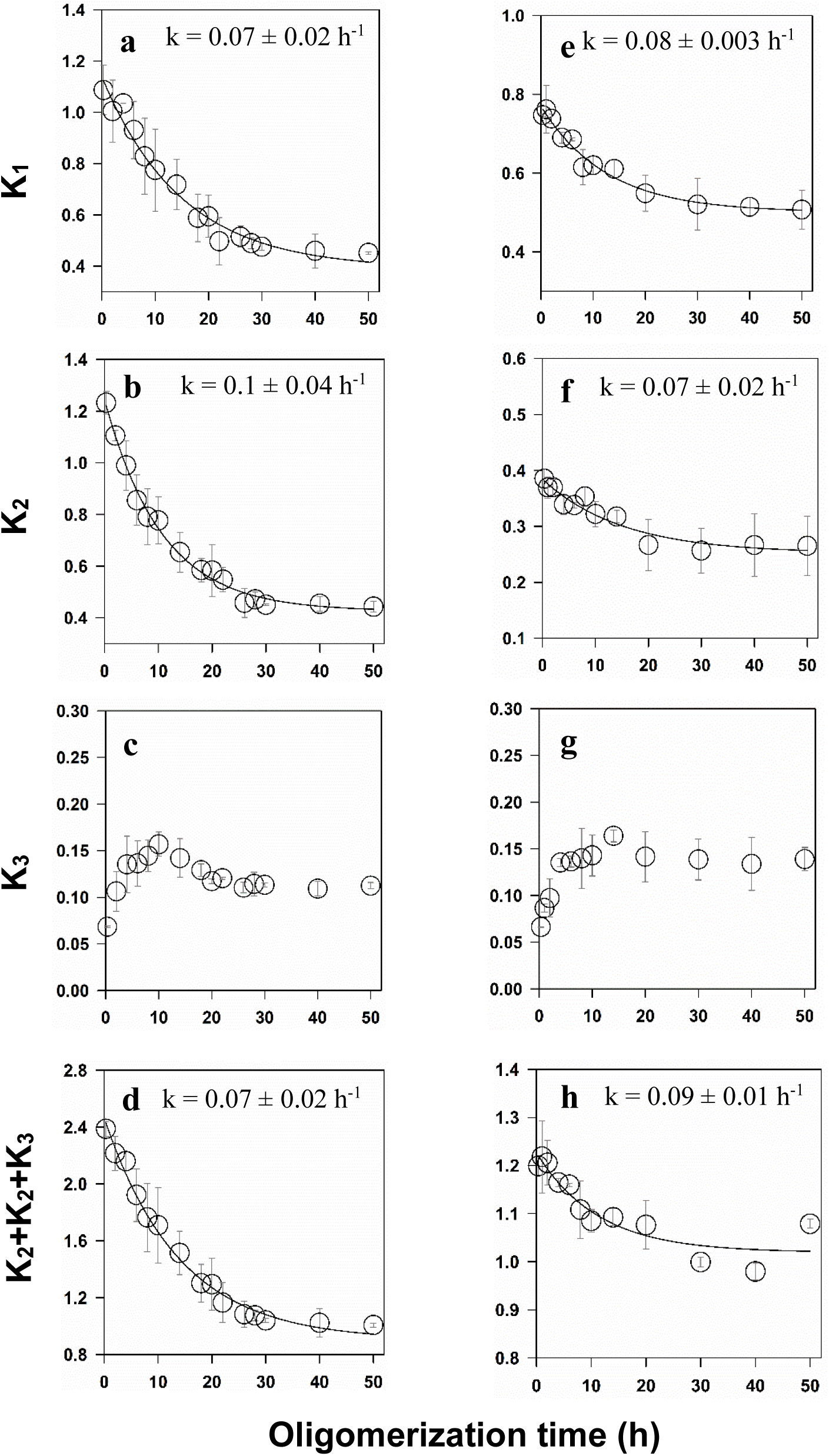
Evolution of the amplitudes of microsecond conformational fluctuations during the oligomerization of moPrP. The evolution of composite amplitudes for the three exponential processes derived from fits to the ACF data is shown as a function of the time of oligomerization probed by the W144/C199-Atto probe (a-d) and the W171/C225-Atto probe (e-h). (a,e) Evolution of K_1_, (b,f) Evolution of K_2_, (c,g) Evolution of K_3_, (d,h) Evolution of the sum of amplitudes (K_1_ + K_2_ + K_3_) showing the global reduction in the amplitude of microsecond fluctuations. The solid lines through the data in panels a, b, d, e, f and h are fits to a single exponential equation, and the values of the apparent rate constants are given in the panels. In each panel, the error bars represents the standard deviations obtained from two independent experiments.

## Discussion

### Use of FCS and PET-FCS to study protein oligomerization

FCS has proven to be a useful tool for studying the oligomerization of proteins including disease-linked proteins such as Aβ (Mittag et al., 2014; Wennmalm et al., 2015), α-synuclein (Takahashi et al., 2007; Nath et al., 2010, 2012; Fan et al., 2022), and a huntingtin fragment (Sahoo, et al., 2016; Sahoo, Drombosky, et al., 2016; Mittag et al., 2019) as well as p53 (Rajagopalan et al., 2011), Rubisco activase (Chakraborty et al., 2012; Kuriata et al., 2014; Serban et al., 2018) and tubulin (Boukari et al., 2003; Krouglova et al., 2004; Sánchez et al., 2004). It has been utilized to study binding reactions such as the binding of SDS (Sen et al., 2021) and tubulin (Elbaum-Garfinkle et al., 2014; Li et al., 2015; McKibben & Rhoades, 2019) to tau. In these studies, as in the present study, multiple protein forms differing in size were present together. In oligomerization studies, the ACF curves shift to longer times, and it is invariably seen, as in the current study, that the ACF curves can be fit adequately to an equation for a single diffusing species (Krichevsky & Bonnet, 2002). The different forms present do not differ in size sufficiently that their diffusion times can be resolved. The fit yields a diffusion time, and hence, a diffusion coefficient that are brightness-weighted averages of the diffusion times and diffusion coefficients of monomer and all oligomeric forms present. In the current study, the progress of the oligomerization reaction was monitored by determination of the apparent diffusion time, τ_D_, at different times of oligomerization. Despite τ_D_ not being proportional to oligomer concentration, the time course of its change nevertheless yielded an apparent rate constant of oligomerization that matched the rate constant of misfolding as measured by far-UV CD measurements (Figure 2).

### MEM analysis reveals oligomer heterogeneity

The MEM analysis of the diffusion component in the ACF makes no prior assumption about the number of diffusing species, and provides a distribution of the diffusion times. The inability of the MEM analysis to resolve between the different species present at any time of oligomerization, is presumably because their hydrodynamic radii are less than 10-fold different from each other.

The shift in the unimodal MEM distribution to longer diffusion times, as well as its broadening, (Figure 5b and e), could indicate a progressive increase in the size of the oligomer population with time, or a fractional increase in the oligomer populations at the expense of the monomer population (Figure S2b). Unfortunately, it is not possible to conclude whether the shift observed for the MEM distribution occurs by one or both of these possible ways. Nevertheless, the observation that the MEM distributions obtained at different times of oligomerization intersect each other at multiple τ_D_ values suggests that growth of the oligomers occurs in a complex manner. It is, however, important to note that the observation that the shift of the peak of the MEM distribution to larger diffusion times occurs with the same apparent rate constant as that of the far-UV CD change indicates not just that the MEM analysis is valid but that the MEM distributions correctly represent how oligomerization proceeds with time.

### Sizes of identified oligomers at pH 4

The mechanism of formation of β-rich oligomer at pH 4 has been extensively studied, and much insight has been obtained (Khan et al., 2010; Singh & Udgaonkar, 2015b; Sabareesan & Udgaonkar, 2016; Sengupta et al., 2017; Sengupta & Udgaonkar, 2019). Nevertheless, while it has been known for a long time that both small and large oligomer populations are formed, the translational sizes of the oligomers were not known. In this study, by measurement of the diffusion times, it was possible to determine the hydrodynamic radii of the small and large oligomers (Figure 3 and Table 1). The hydrodynamic radii can be used to estimate the number of monomeric units in O_S_ as well as in O_L_. With the assumption that the oligomers are spherical with constant specific volume, the number, n, of monomeric units in an oligomer is given by n = (R_h_^O^/R_h_^M^)^3^, where R_h_^M^ and R_h_^O^ are the hydrodynamic radii of the monomer and oligomer, respectively (Chakraborty et al., 2012; Nath et al., 2012). Thus, the number of monomeric units in O_S_ and O_L_ can be estimated to be 15 and 55, respectively. Previous estimates of the sizes of O_S_ and O_L_ at pH 2, indicated that they were comprised of 12 and 50 monomeric units, respectively (Singh et al., 2012). It should be noted, however, that the oligomers formed at pH 2 are known to be different from those formed at pH 4, in that, unlike the latter, they go on to form worm-like fibrils (Jain & Udgaonkar, 2011).

### Persistence of microsecond dynamics in the oligomers

The observation that the fluctuations persist during the course of oligomerization, with their composite time constant not changing and only their composite amplitudes reducing, was surprising. The fluctuations are seen in the purified oligomers too (Figure 3 and Table 1), indicating that the fluctuations seen at the end of the oligomerization reaction could not have originated from the ∼8 % residual monomer that remains. The oligomers are both rich in β-sheet structure, while the monomer is rich in α-helical structure, and the persistence of the fluctuations in the oligomers on similar timescales suggests that monomer, as well as and monomer present in oligomers, have similar flexibility. Indeed, HX-MS studies on oligomer and monomer suggest that the flexibilities of most, but not all (see later) of the sequence segments are not very different (Singh & Udgaonkar, 2015b; Sabareesan & Udgaonkar, 2016). It should be noted, however, that the timescales of the fluctuations cannot be determined in the HX-MS experiments.

The timescale of contact formation between the Trp donor and the Atto-labelled Cys is not determined only by the distance between the two sites, but also by the friction experienced during local segmental motion. This friction can have both solvent-dependent and solvent-independent contributions. The solvent-dependent contribution is related to solution viscosity, which affects diffusive segmental motion and contact formation (Waldauer et al., 2010; Sherman & Haran, 2011). The solvent-independent contribution, or internal friction, can arise from intrachain interactions, steric constraints, dihedral-angle barriers and roughness of the conformational energy landscape (Neuweiler et al., 2009; Soranno et al., 2017). Thus, local interactions in the protein do not merely change the probability of contact formation; they can also modify the internal friction that governs how rapidly such contacts form and dissociate. In this context, the persistence of similar relaxation times suggests that oligomerization does not greatly change the effective frictional barrier for the local motions monitored by the two PET pairs. The decrease in the amplitudes, however, indicates that PET-quenched dark states, N* and N**, are sampled less in the oligomeric state.

It is possible that the likelihood of native-state fluctuations that bring the Trp donor in contact with the Atto moiety is dictated more by the sequence separation between the Trp residue and the Cys residue with the Atto adduct. If the monomer and the monomeric unit in oligomer have similarly flexible structures, then the probability of two residues coming into contact might be dictated by the number of residues in the sequence separating them, as is the case for random polymers and loops in proteins (Szabo et al., 1980; Pastor et al., 1996; Lapidus et al., 2000; Bhatia & Udgaonkar, 2022). The observation that the amplitudes of the fluctuations, K_1_ and K_2_, are lower in the oligomers, while the time constants are invariant, indicates that the rate constant of association is slower in the oligomers and the rate constant of dissociation of the PET complex is faster. This indicates that fluctuations between the Trp residue and the Atto moiety, which bring them into contact, are less likely in the oligomer than in the monomer, although not very significantly so. Hence, oligomerization mainly reduces the population of the PET-quenched dark states, N* and N** rather than eliminating the local motions.

In FCS measurements of Atto655–Trp contact dynamics in the denatured B1 domain of protein L too, the amplitude of the fast contact-dependent process was found to decrease with increasing GdnHCl concentration, whereas its lifetime remained nearly independent of denaturant concentration, consistent with a reduced probability of dye–Trp contact during chain expansion (Sherman & Haran, 2011). That study, however, contradicts another study of the unfolded B1 domain of protein L, which showed that dilution from high GdnHCl causes chain compaction and a 100-to 500-fold decrease in the intramolecular diffusion coefficient, indicating that contact dynamics can slow markedly as the chain becomes more compact (Waldauer et al., 2010). Single-molecule studies on unfolded proteins have further shown that the reconfiguration time, τ_r_, is sensitive to denaturant-dependent chain expansion or collapse and to internal friction (Nettels et al., 2007; Borgia et al., 2012; Soranno et al., 2012, 2017). In studies of SDS-induced binding and folding of tau-K18, PET-FCS has revealed that local conformational fluctuations persisted in both the disordered U state and the SDS-bound helical FL5 state, indicating that structure acquisition did not abolish fast local dynamics. In that case, however, the fast relaxation time increased from 200–400 ns in U to 800–1200 ns in FL5, whereas the amplitude showed only marginal dependence on SDS concentration (Sen et al., 2021).

### Dynamics as a reporter for conformational conversion

The observation that the likelihood of contact formation at either site in the monomeric protein decreases concurrently with conformational conversion during oligomer formation, suggests that the changes in the dynamics are reporting on local structural changes. The observation that changes at the α1-α3 and α2-α3 interfaces occur concurrently (Figure 4) suggests that the local structure changes at the two interfaces occur simultaneously during the process of conformational conversion during oligomerization. In this context, it should be noted that the addition of 150 mM NaCl, which triggers the process of oligomerization at pH 4, has similar effects on the timescales of fluctuations at the two sites in the monomer, although the effect is slightly more at the α2-α3 interface.

### Dynamics at the α1–α3 interface is affected slightly more strongly than in the α2–α3 core

It is interesting to note that the fractional decrease of the total change in the amplitudes of the fluctuations is more for those monitored by the W144/C199-Atto PET pair than those monitored by the W171/C225-Atto probe. Unfortunately, little is known about the structure of the protein in the oligomeric state. Earlier FRET studies (Sengupta & Udgaonkar, 2019) had indicated that, during oligomerization, the fractional increase in the distance separating residues 144 and 199 is more than that in the distance separating residues 197 and 223. This would account for the larger fractional decrease in the amplitude of fluctuations between Trp144 and Cys199-Atto than for the amplitude of fluctuations between Trp171 and Cys225-Atto. These site-specific FRET measurements also showed separation of the β1–α1–β2 subdomain from the α2–α3 subdomain during oligomer formation (Sengupta & Udgaonkar, 2019). HX-MS studies (Singh & Udgaonkar, 2015b) have shown that the sequence stretch separating Trp171 from Cys225-Atto in the oligomer acquires more stable structure during the transition from monomer to oligomer than does the sequence stretch separating Trp144 from Cys199 in the oligomer. That result is consistent with the dynamics between Trp171 and Cys225-Atto undergoing a smaller fractional decrease in amplitude than that between Trp144 and Cys199-Atto. HX-MS studies also showed that the segment that was α1 in the native state becomes more solvent-exposed in the oligomeric state, whereas segments that comprise the erstwhile α2–α3 region in N become incorporated into the protected oligomer core (Singh & Udgaonkar, 2015b). Thus, the large fractional decrease in the PET-FCS amplitude at what was the α1–α3 interface in N is consistent with a larger structural rearrangement across this interface. In contrast, the smaller fractional change seen at what was the α2–α3 interface may be because this region becomes part of the protected oligomer core.

## Conclusion

This PET-FCS study of the oligomerization of moPrP provides a characterization of not only the evolution of oligomer size, but also the evolution of microsecond dynamics across two regions, corresponding to the erstwhile α1-α3 and α2-α3 interfaces present in the native protein. The sizes of the small and large oligomers, O_S_ and O_L_, that are formed, have been determined and the distinctive dynamics at the two erstwhile interfaces have been characterized. The fluctuations that are observed in the native state ensemble appear to persist in O_S_ and O_L,_ occur on similar timescales, but are damped to different extents. The differences in the damping of fluctuations across the two regions are shown to be correlated with the distinct structural changes known to occur there during the course of oligomerization. Hence, the observation that the changes in dynamics across the two regions occur concurrently suggests that the structural changes also occur together.

## Materials and methods

### Buffers and reagents

All chemicals used were of the highest purity grade and procured from Sigma unless otherwise mentioned. Guanidine hydrochloride (GdnHCl) was purchased from Himedia. Experiments were carried out in 10 mM sodium acetate buffer, pH 4, and aggregation buffer contained, in addition, 150 mM NaCl. Buffers used for the FCS experiments contained 0.05% Tween 20 to prevent protein adsorption onto the surfaces of the coverslips.

### Protein expression, purification and site-specific labelling

Single-Trp as well as single-Cys-containing moPrP variants were expressed in *Escherichia coli* BL21(DE3) codon plus cells and purified from inclusion bodies using previously described procedures (Sengupta & Udgaonkar, 2017). Following Ni-NTA affinity purification under denaturing conditions, the proteins were refolded by stepwise removal of GdnHCl in the presence of GSH/GSSG. The refolded proteins were further purified using a HiTrap CM FF weak cation-exchange column, dialysed against Milli-Q water, flash-frozen, and stored at −80°C.

Purified protein was reduced with 10X TCEP (Tris (2-carboxyethyl) phosphine hydrochloride) in 10 mM sodium acetate, at pH 4.0 and 4 °C for 12 h, residual GSH and TCEP were removed, followed by immediate reaction with a 10-fold excess of Atto 655 maleimide in 20 mM Tris-HCl pH 7.8 for 12 h at 4 °C in the dark. Excess dye was removed using a HiTrap desalting column (Cytiva). Each protein was identified by electrospray ionization mass spectrometry, and protein concentrations were determined by absorbance measurements, using an ε value 22,450 M^-1^cm^-1^ at 280 nm for the unlabelled protein, and using an ε value 1,25,000 M^-1^cm^-1^ at 663 nm for Atto-labelled protein (Goluguri et al., 2019).

### Oligomerization monitored by far-UV circular dichroism (CD) spectroscopy

Aggregation experiments were initiated by diluting purified monomer two-fold into 2x-aggregation buffer so that it was finally in 10 mM sodium acetate, 150 mM NaCl, pH 4 at 37 °C. The final protein concentration was 100 µM. 10 nM of labelled protein was added as the dopant, so that the protein used for the CD experiment was the same as that used for PET-FCS measurements. Samples were incubated at 37 °C without agitation. To monitor misfolding kinetics, aliquots were withdrawn at different times and probed by far-UV CD measured at 222 nm at 25 °C. The CD signal was converted to “fraction misfolded” by normalizing between values of 0 for the native monomer and 1 for the far-UV CD signal at the end of the oligomerization reaction. Kinetic traces (fraction *vs* time) were fit to a single exponential equation (Equation 1) to obtain apparent rate constant of oligomerization:

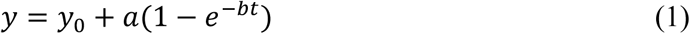

### Oligomerization monitored by PET-FCS

PET-FCS measurements were carried out on a MicroTime 200 time-resolved confocal microscope (PicoQuant, Germany), as described previously (Goluguri et al., 2019). Atto 655 was excited using a 637 nm pulsed diode laser operating at 40 MHz with an excitation power of 30 µW, chosen to minimize triplet-state population and photobleaching. The excitation beam was focused into the sample using a 60×, NA 1.2 water-immersion objective, and fluorescence was collected through the same objective in the epifluorescence mode using a 480/645 dual-band dichroic mirror. The emitted fluorescence was passed through a 690/ 70 nm band-pass filter and a 50 µm pinhole, split using a 50:50 beam splitter, and detected by two single-photon avalanche photodiodes. The photon streams from the two detectors were cross-correlated to eliminate detector-afterpulsing artifacts. The confocal observation volume was calibrated in each experiment by measuring the autocorrelation functions (ACF) of free Atto 655 under identical optical conditions. To exclude contributions from excitation power-dependent fluorophore photophysics, control measurements were carried out at excitation powers ranging from 15 to 45 µW. The amplitudes and time constants of the exponential components were independent of excitation power. To assess contributions from residual free Atto 655, labelled samples were additionally buffer-exchanged using a 10 kDa Amicon centrifugal filter, resulting in a further approximately 10^6^-fold dilution of unbound dye. The exponential components remained unchanged after buffer exchange, indicating that exponential components did not arise from residual free dye.

For monitoring oligomerization, 100 µM Trp-less moPrP was doped with 10 nM Atto 655 labelled protein incubated in 10 mM sodium acetate buffer containing 150 mM NaCl at pH 4.0 and 37 °C. The dopant was either W144/C199-Atto moPrP or W171/C225-Atto moPrP. PET-FCS measurements were taken at different times during the oligomerization process. Coverslips were passivated with a coating of 0.1 % BSA. 0.05% Tween 20 was included in the buffer to minimize non-specific adsorption. For each time of oligomerization, the ACFs were collected for 15–40 min to ensure good quality data. The ACFs were analyzed to obtain the diffusion time and the individual fluctuation times, as described below.

### Data Analysis for fitting of the ACFs

Photon arrival time records from each measurement were autocorrelated (using SymphoTime-64, PicoQuant) to obtain the fluorescence intensity autocorrelation function (ACF). The experimental ACFs were then fitted to a three-dimensional diffusion term plus three independent single-exponential decay terms, using Equation 2:

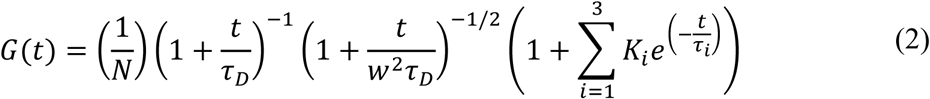

τ_D_ is the diffusion time, N is the mean number of particles in the confocal volume, and K_i_ and τ_i_ are the amplitude (also equilibrium constant) and characteristic decay time of the i^th^ relaxation process (Goluguri et al., 2019). The fit was performed using SigmaPlot, which allowed a robust extraction of amplitudes (K_1_, K_2_, K_3_) and time constants (τ_1_, τ_2_, τ_3_) of all three sub-millisecond relaxation processes. *w* is the aspect ratio which is the ratio of the radial to axial dimensions of the confocal volume.

### Size-exclusion chromatography purification of oligomers

100 µM Trp-less moPrP doped with 10 nM W144/C199-Atto moPrP or W171/C225-Atto moPrP was incubated in aggregation buffer at 37 °C. At completion of oligomerization, the reaction mixture was loaded onto a Bio-Sil SEC 400-5 column equilibrated with aggregation buffer. The monomer, small oligomer, and large oligomer fractions were collected separately based on their elution profiles.

The isolated SEC fractions were used immediately for PET-FCS measurements under the same aggregation buffer conditions. ACFs were recorded for the monomer, small oligomer, and large oligomer fractions of samples doped with both W144/C199-Atto moPrP and W171/C225-Atto moPrP. The ACFs were fitted using Equation 2. The diffusion time, τ_D_, that was obtained for the monomer and each oligomer species was used to determine the hydrodynamic size of the separated species, using the Stokes–Einstein equation:

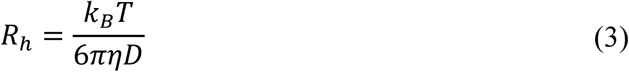

where *k_B_* is the Boltzmann constant, *T* is the absolute temperature, and *η* is the viscosity of the solution. *D* is diffusion coefficient which is equal to 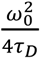 where *ω_0_* is the lateral radius of the confocal observation volume. To determine R_h_ of each species, the τ_D_ value for the free dye was obtained under identical conditions of measurement. Since the R_h_ value of the dye was known to be 0.6 nm, the R_h_ value of the protein species could be determined.

Both oligomers are assumed to be spherical in shape; the measured Rₕ values were used to estimate the number of monomeric units in O_S_ and O_L_. The number of monomeric units, *n*, in the small and large oligomers was estimated (Chakraborty et al., 2012) using:

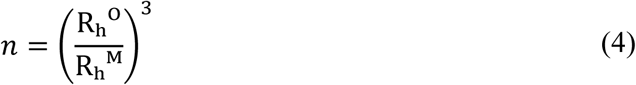

Rₕᴼ is the mean hydrodynamic radius of either O_S_ or O_L_, averaged across the two PET probes, and Rₕᴹ is the mean hydrodynamic radius of the monomer measured under the same conditions.

### Analysis of the ACF using the Maximum Entropy Method (MEM)

During the oligomerization reaction, the fluorescently labelled moPrP molecules are present in a heterogeneous mixture of monomeric and oligomeric species. Conventional analysis of the diffusion component requires the number of diffusing species to be specified before fitting, which may not be appropriate when the sample contains a distribution of oligomeric sizes. Hence, the diffusion component of the ACFs was additionally analyzed using the Maximum Entropy Method FCS (MEMFCS), as described previously (Sengupta et al., 2003).

In MEMFCS analysis, the diffusion component of the ACF is represented as a weighted sum of multiple diffusion components:

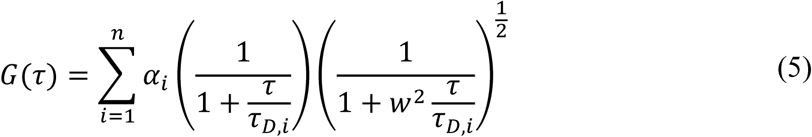

where *α_i_* is the amplitude of the *i*^th^ diffusion component, *τ_D_*_,*i*_ is the corresponding diffusion time, *n* is the number of diffusion components used in the analysis, and *w* is the ratio of the radial to axial dimensions of the confocal observation volume. In this analysis, the diffusion times were kept fixed over a defined range, while the amplitudes were varied to obtain the diffusion-time distribution that best described the experimental ACF.

The amplitudes were optimized by minimizing the deviation between the calculated and experimental ACFs while maximizing the entropy, S, of the distribution:

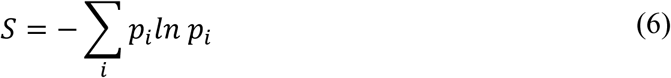

where 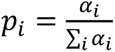. This procedure allows the ACF to be described as a distribution of diffusion times, rather than as one or a few discrete diffusing species. Thus, MEMFCS is a useful way to examine diffusional heterogeneity in samples where multiple species are present but their diffusion times are not sufficiently separated to be resolved reliably by conventional discrete-component fitting.

For the moPrP oligomerization reactions, MEMFCS was applied to ACFs collected at different reaction times from samples doped with either W144/C199-Atto moPrP or W171/C225-Atto moPrP. The peak of the diffusion time distribution was used as the representative diffusion time at each time point.

## Acknowledgements

I thank Prof. Jayant B. Udgaonkar for his supervision, scientific guidance, and valuable suggestions and corrections to the manuscript. I thank Dr. Jeetender Chugh for valuable discussions, critical reading, and suggestions on the manuscript. I thank my laboratory members and Dr. Sarita Puri for helpful comments. I thank Rama Reddy Goluguri and Dr. Ishita Sengupta for generating the PET-pair mutant variants. I thank Vicky Vishvakarma from Prof. Sudipta Maiti’s laboratory for his help in learning MEMFCS analysis. I thank the University Grants Commission (UGC) for a graduate research fellowship. This work was supported by a National Science Chair grant from the Anusandhan National Research Foundation to Prof. Jayant B. Udgaonkar.

## SUPPLEMENTARY INFORMATION

**Figure S1.**
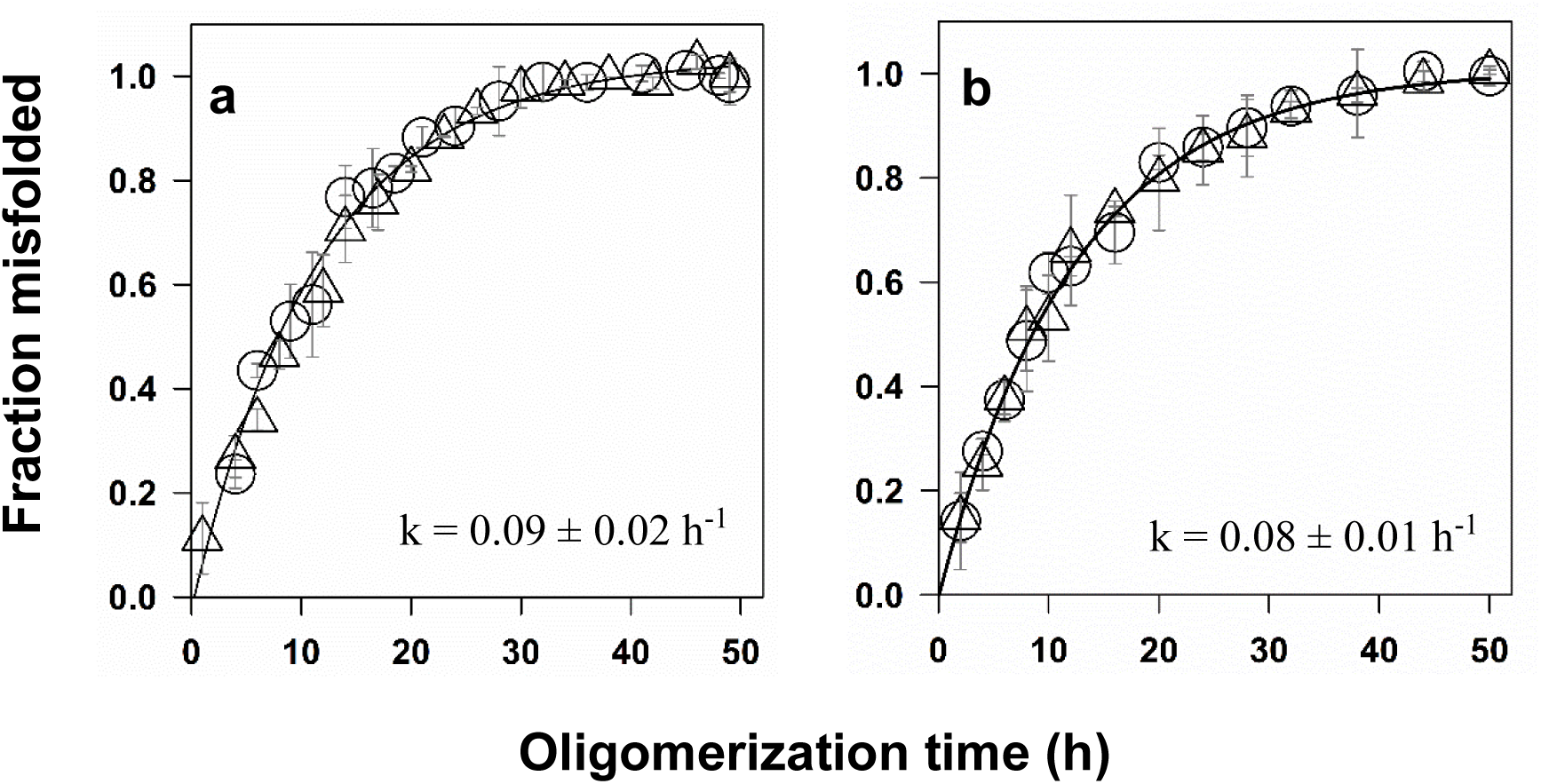
Oligomerization kinetics of Trp-less moPrP doped with labelled moPrP in the presence of Tween 20. The oligomerization reactions of 100 µM Trp-less moPrP, doped with. (a) 10 nM W144C199-Atto moPrP and (b) 10 nM W171C225-Atto moPrP were monitored by far UV CD at 222 nm, in 150 mM NaCl, 10 mM NaOAc, pH 4 at 37 °C. The kinetics were determined in the presence (ο) and absence (Δ) of 0.05% Tween 20. The solid lines represent fits of the data to a single exponential equation (Equation 1). The error bars represent the standard deviations of the mean determined from two independent measurements.

**Figure S2.**
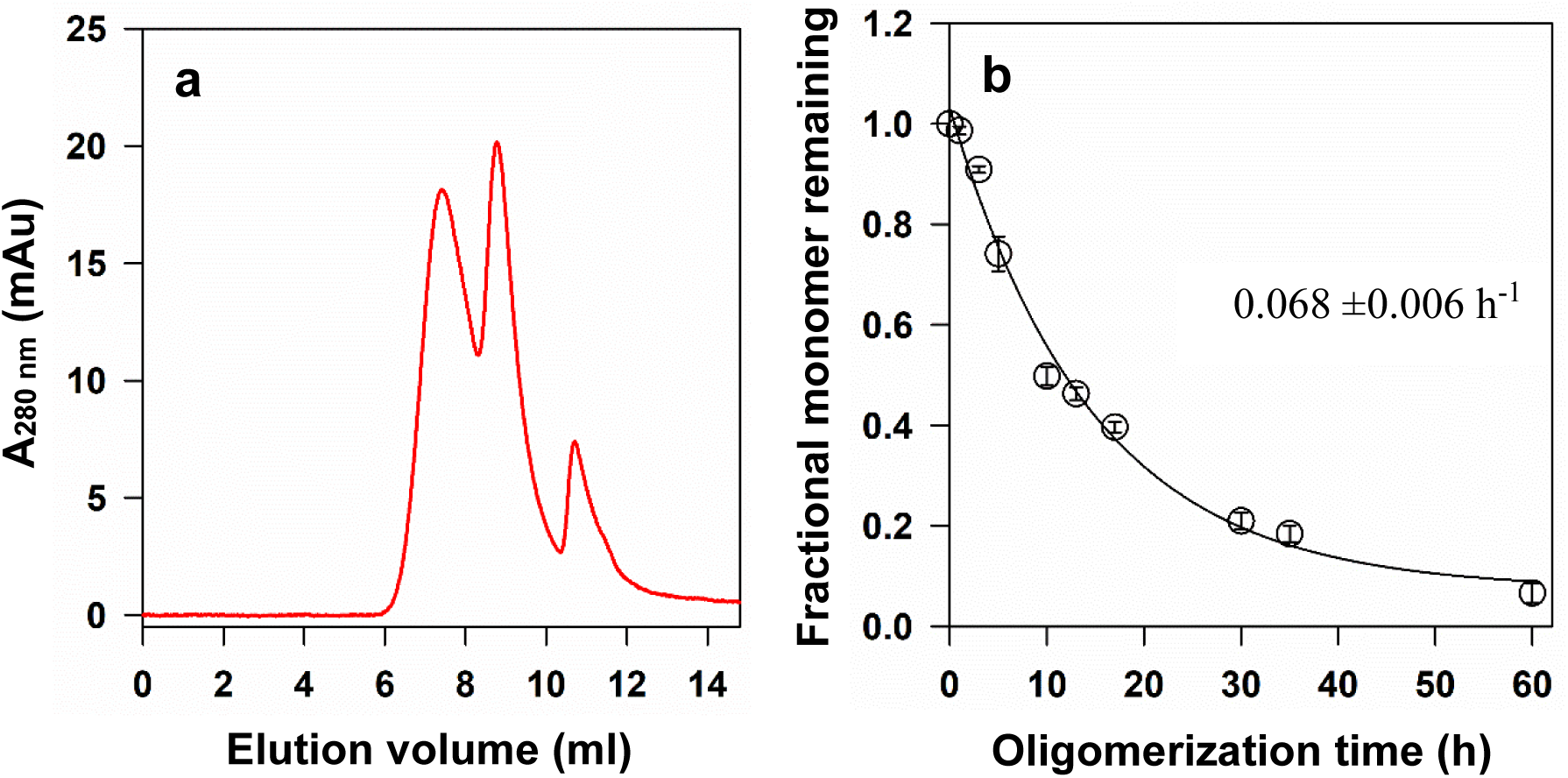
SEC analysis of monomer loss during oligomerization of moPrP. (a) SEC profile of the reaction mixture at the end of oligomerization, showing the distribution of protein as monomer (M), small oligomer (O_S_), and large oligomer (O_L_). (b) Time-dependent decrease in monomer concentration during oligomerization. The error bars in panel b represent the standard deviations obtained from two independent experiments.

**Figure S3.**
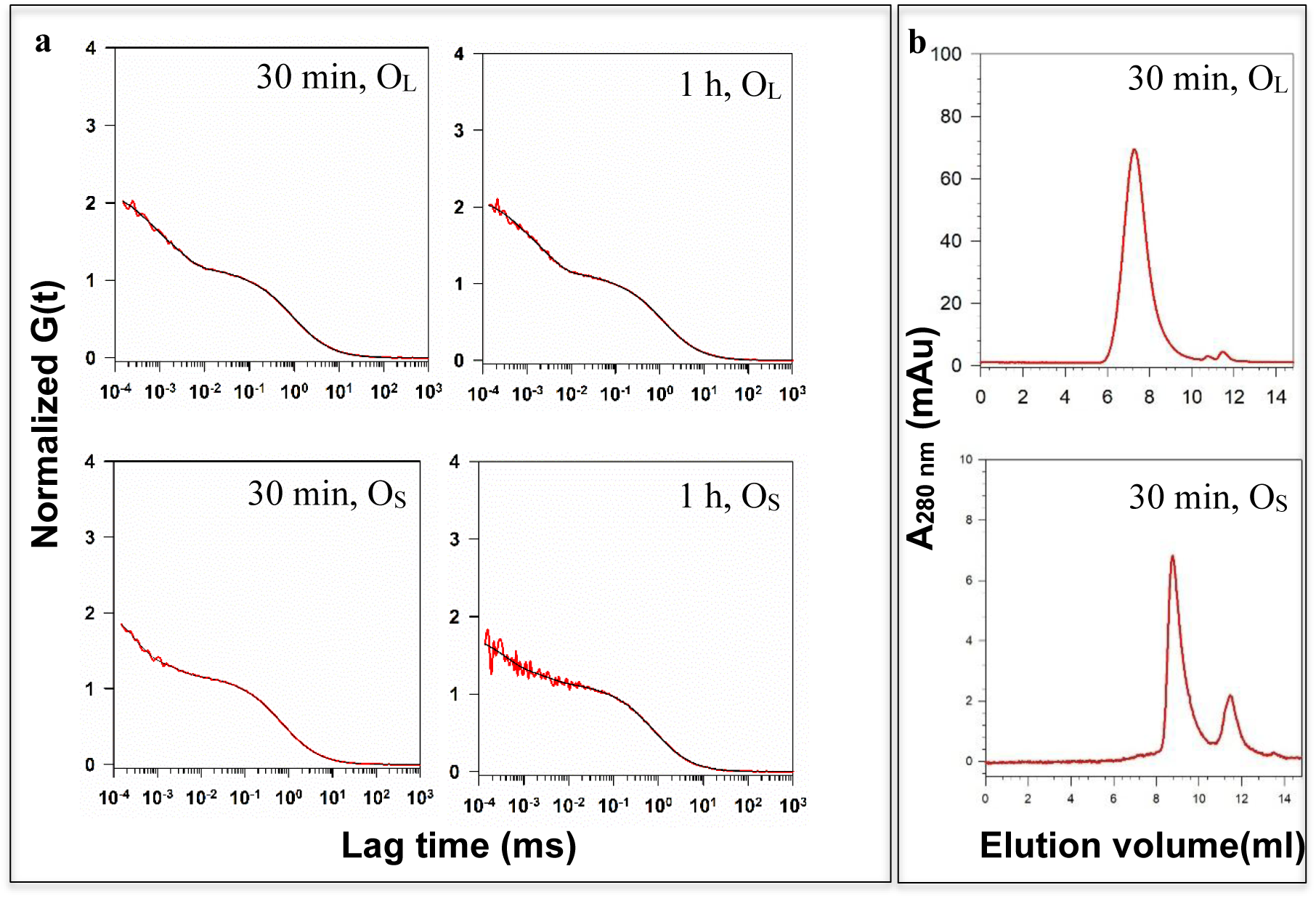
Purified oligomers remain pure over the timescale of PET-FCS measurements. (a) ACFs of oligomers of W144/C199-Atto moPrP at different times after purification. (b) SEC of oligomers of W144/C199-Atto moPrP at 30 min after purification.

**Table S1.** Parameters describing the ACFs shown in Figure S3.

|  | **30 min, O<sub>L</sub>** | **1h, O<sub>L</sub>** | **30 min, O<sub>s</sub>** | **1 h, O<sub>s</sub>** |
| --- | --- | --- | --- | --- |
| **K<sub>1</sub>** | 0.4 | 0.33 | 0.65 | 0.38 |
| **τ<sub>1</sub> (μs)** | 0.41 | 0.46 | 0.28 | 0.36 |
| **K<sub>2</sub>** | 0.54 | 0.60 | 0.24 | 0.20 |
| **τ<sub>2</sub> (μs)** | 2.8 | 3 | 2.5 | 3.1 |
| **K<sub>3</sub>** | 0.08 | 0.09 | 0.07 | 0.06 |
| **τ<sub>3</sub> (μs)** | 51 | 56 | 49 | 33 |
| **τ<sub>D</sub> (μs)** | 929 | 1164 | 705 | 770 |

**Figure S4.**
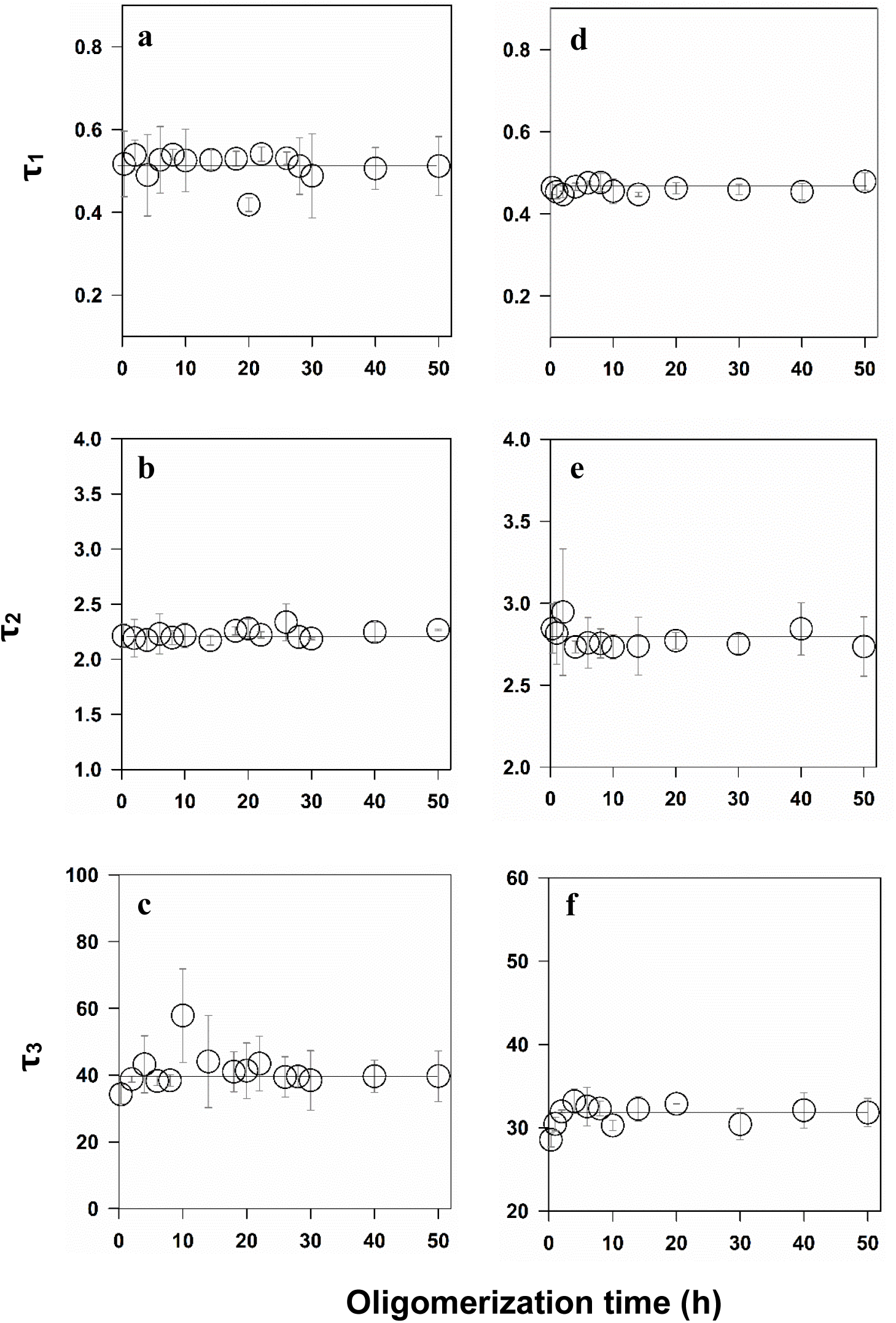
Evolution of time constants of microsecond configuration fluctuation during oligomerization. The evolution of the three time constants (τ_1_, τ_2_, τ_3_) obtained from the fits of the data in Figure 6 to Equation 2 is shown as a function of oligomerization. (a-c): W144/C199-Atto probe and (d-f): W171/C225-Atto probe. (a,d) τ_1_ evolution (b,e) τ_2_ evolution (c,f) τ_3_ evolution. The error bars represent the standard deviations of the mean determined from two independent measurements.

